# An untargeted metabolomics analysis of exogenous chemicals in human milk and transfer to the infant

**DOI:** 10.1101/2022.03.31.486633

**Authors:** Sydney Thomas, Julia M. Gauglitz, Anupriya Tripathi, Fernando Vargas, Kerri Bertrand, Jae H. Kim, Christina Chambers, Pieter C. Dorrestein, Shirley Tsunoda

## Abstract

Human milk is the optimal infant nutrition. However, while human-derived metabolites such as lipids and oligosaccharides in human milk are regularly reported, the presence of exogenous chemicals (such as drugs, food, and synthetic compounds) are often not addressed. To understand the types of exogenous compounds that might be present, human milk (n=996) was analyzed by untargeted metabolomics. This analysis revealed that lifestyle molecules such as medications and their metabolites, food, industrial sources such as plasticizers, cosmetics, microbial molecules, and other personal care products are found in human milk. We provide further evidence that some of these lifestyle molecules are also detectable in the newborn’s stool. Thus, this study gives important insight into the types of exposures infants receiving human milk might ingest due to the lifestyle choices, exposure, or medical status of the lactating parent.

## Introduction

Human milk is the gold standard of infant nutrition, supplying a complete diet including micronutrients, macronutrients, bioactive compounds, immunomodulatory components, and hormones.^1^ In the US, over 80% of infants receive human milk for varying duration, with the American Academy of Pediatrics recommending breastfeeding for the first 6-12 months of life.^2^ Many studies have focused on biological metabolites produced by the lactating parent, but human milk also contains chemicals derived from exogenous sources, such as personal care products, food, or drugs.^3–6^ These lifestyle molecules have the same potential to shape the developing infant as human-derived compounds.^7,8^ To better understand the exogenous molecules that are present in human milk, we analyzed six publicly-available untargeted metabolomics datasets that included 996 human milk samples. Our results uncover a large range of lifestyle metabolites, including several medications and synthetic compounds from personal care products. We further show with paired mother-infant samples that several of these molecules are transferred from human milk to the infant in detectable quantities. These data provide the first step towards understanding the full range of compounds that may be passed from the lactating parent to infant via human milk.

## Methods

We analyzed six datasets comprising 996 human milk samples collected over a three year period from Mommy’s Milk, a Human Milk Research Biorepository.^9^ All women provided written consent for use of samples under UCSD IRB numbers 130658 and 151713. Details for each dataset are included in **Supplementary Table 1**. Datasets are publicly available in the MassIVE database (http://massive.ucsd.edu).

Human milk samples were extracted using multiple solvent concentrations: 80:20 MeOH:water, 50:50 MeOH:water, 80:20 EthOH:water, and 80:20 EthOH:water. Untargeted metabolomics was performed using an UltiMate 3000 liquid chromatography system (Thermo Scientific) coupled to a Maxis Q-TOF (Bruker Daltonics) mass spectrometer with a Kinetex C18 column (Phenomenex). Samples were run using a linear gradient of mobile phase A (water 0.1% formic acid (v/v)) and phase B (acetonitrile 0.1% formic acid (v/v)). A representative linear gradient consisted of 0-0.5 min isocratic at 5% B, 0.5-8.5 min 100% B, 8.5-11 min isocratic at 100% B, 11-11.5 min 5% B, and 11.5-12 min 5% B. Datasets were run in both positive and negative mode. All solvents used were LC-MS grade.

To gain insight into exposure molecules, a classical molecular network was created both for each dataset separately and all datasets together using GNPS (http://gnps.ucsd.edu).^10,11^ Default settings were used except in these cases: precursor ion mass tolerance and MS/MS fragment ion tolerance were set to 0.02 Da, cosine values were set to 0.6, and matched peaks were set to 5. These settings are appropriate for an exploratory study as they are less stringent and lead to more library matches, but they may also increase false positive identifications (passatutto estimates an FDR of 1.5% using these settings).^12^ All datasets, settings, and analysis are publicly available using the links included in **Supplementary Table 1**.

After molecular networking, networks containing unique drugs and synthetic compounds were identified (**Supplementary Table 2**). Each of these compounds was manually annotated to confirm library annotations. In some cases, exogenous compounds were added as internal standards during sample preparation. In this case, any compounds in the same network were inspected using the GNPS Dashboard to ensure that they were true hits and did not exist in control samples.^13^

## Results

Untargeted metabolomics is a mass spectrometry-based method that can measure thousands of metabolites (compounds <1500 Da) in a single sample. Thus, this method provides a high-throughput, unbiased survey of the chemical components in a biological sample. Despite these advantages, the use of untargeted metabolomics in the study of human milk (and other biological fluids) has been historically limited by the technical challenges of spectral annotation. While most MS/MS experiments produce tens of thousands of spectra which correspond to thousands of unique molecules, the vast majority of these spectra will not match existing spectral libraries. To overcome this obstacle, we employed two cutting-edge tools on six publicly-available untargeted metabolomics datasets. First, we used molecular networking to group spectra by spectral similarity.^11^ This method organizes data into networks where each node represents a different spectra, allowing the user to determine the chemical characteristics of an unannotated spectra by comparing it to annotated nodes in the same network. In addition, we employed a suspect library that annotates spectra based on known chemical modifications (*in preparation*). While these compounds do not have exact library matches, their structures can often be deduced by comparing peaks and chemical composition to a parent compound. Using these methods, we identified roughly 1600 unique spectral matches in six publicly available human milk datasets (**Supplementary Table 1**). The majority of these compounds (70%) were identified in a single dataset, while the rest were identified in multiple datasets. In total, this led to 1121 unique spectral matches between all six datasets, and resulted in at least partial annotation of 23% of the total spectra.

As expected, we identified a large number of known biological components in human milk. This includes a range of amino acids, sugars, vitamins, and fatty acids. However, the dataset also contained a range of exogenous components, including >30 drugs, >25 compounds from food, and >15 synthetic compounds (**Supplementary Table 2**). Of the 33 drugs identified, five were not included the NIH Drugs and Lactation Database (LactMed), including the antibiotics sulfamethazine, sulfadimethoxine, as well as the aspirin metabolite 2,3-dihydroxy-benzoic acid, the antiandrogen flutamide, and the AMPK activator AICAR.^3^ However, the identification of sulfamethazine and sulfadimethoxine should be viewed with caution, as these compounds can be used as internal controls for mass spectrometry. In general, the two largest classes of drugs were antibiotics and antidepressants. In several cases, untargeted metabolomics did not just identify parent compounds for these drugs, but also known degradation products (**Figure 1**). The metabolites related to sulfonamide antibiotics were particularly varied (**Figure 1a**). Specifically, we identified six metabolites of sulfamethazine, two metabolites of sulfapyridine, and one metabolite of sulfadimethoxine. Mirror plots for each sulfonamide metabolite are available in **Supplementary Figure 1**, while chemical information and unique spectral peaks are listed in **Supplementary Table 3**. Although we were able to assign several of these metabolites to known compounds, several more could not be annotated based on public data.^14,15^ Identified metabolites include 2-Acetyl-N-[4-[(4,6-dimethylpyrimidin-2-yl)sulfamoyl]phenyl]benzamide (Sulfamethazine 1, 425.149 m/z), Methyl 4-{[(4-{[(4,6-dimethylpyrimidin-2-yl)amino]sulfonyl}phenyl)amino]carbonyl}benzoate (Sulfamethazine 2, 441.143 m/z), and Acetylsulfapyridine (Sulfapyridine 2, 292.075 m/z). While it is possible that these metabolites are inactive, the presence of known active drug metabolites, such as nortriptyline or amitriptyline-*N*-oxide, suggests that multiple active forms of a drug could be transferred to the infant through human milk (**Figure 1b**). In fact, in cases where known drug metabolites were identified in human milk datasets, the drug metabolites occur with similar frequency to their parent compounds. For example, descalinose azithromycin (34 samples) occurred in 75% as many samples as its parent compound azithromycin (44 samples). The frequency and variety of these drug metabolites suggests that variable formation of these compounds could impact the therapeutic or toxic effects in infants. In addition to drug compounds, we also identified a range of bacterial quorum signaling molecules in human milk (**Figure 1d**). This highlights the effects of the human milk microbiome on human milk metabolism, and raises the possibility that metabolism of drugs in human milk may not always follow known human drug metabolism pathways.

**Figure 1:**
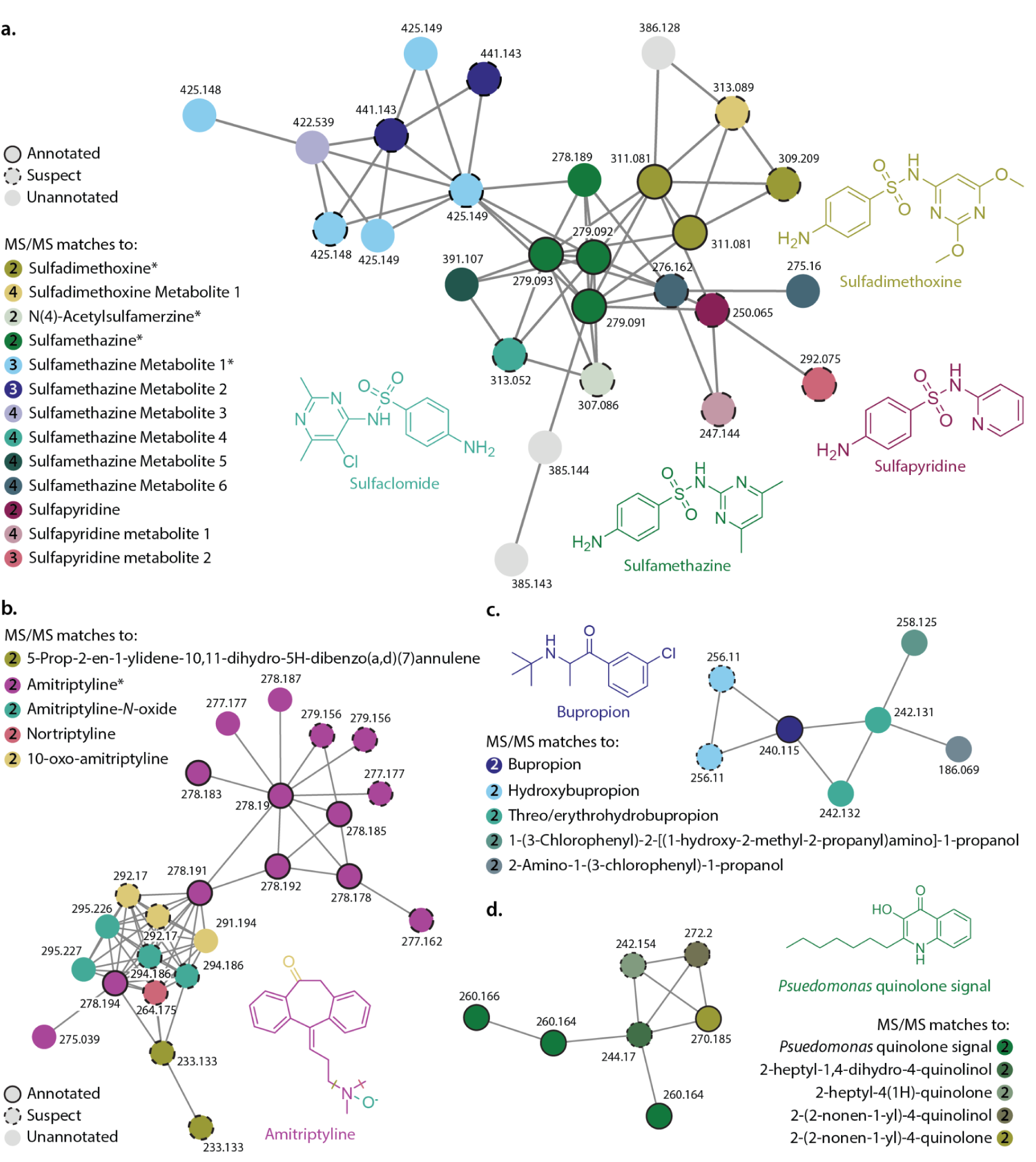
Molecular networks containing drug and food metabolites in human milk. **a**. Molecular network containing sulfonamide antibiotics and antibiotic metabolites. **b**. Molecular network containing antidepressant amitriptyline and multiple known degradation products. **c**. Molecular network containing the antidepressant bupropion and its metabolites. **d**. Molecular network containing bacterial quorum sensor Pseudomonas quinolone signal and related metabolites. Network nodes with full annotations from the original analysis are shown with black outlines, suspect annotations are shown with dotted outlines, and unannotated nodes have no outline. Asterisks (*) mark compounds that were identified in solvent controls in at least one dataset, number in circle indicates MS/MS match level.^16^ All colored nodes in these networks have been manually annotated by spectral comparison to library compounds and other related spectra. Library and spectral m/z, library quality, m/z error, and shared peaks for each MS/MS match are included in **Supplementary Table 2**.

To further understand whether drugs are directly transferred to the infant, we used a publicly-available dataset containing paired metabolomic samples from 42 mother-infant dyads. Samples included the lactating parent’s human milk and infant oral, skin, and stool samples. In total, 431 metabolites occurred in both the parent and infant samples in at least one dyad. The most common paired metabolites included human-derived compounds such as fats, sugars, and amino acids, but several exogenous compounds were also identified. For example, while not all drugs in our larger human milk analysis were found in this smaller cohort, we did identify the antibiotics erythromycin, azithromycin, and clindamycin in paired samples (**Figure 2**). In these samples, the antibiotics were most commonly present in both human milk and infant stool, suggesting that antibiotics in human milk can traverse the entire intestinal tract of the infant. Besides antibiotics, we were also able to identify the beta blocker labetalol and the pain reliever acetaminophen in at least one of the dyads (**Figure 2b**). Acetaminophen in particular was identified in more infant than parent samples, likely due to direct administration to the infant. In addition to known compounds, several unannotated compounds showed remarkably similar patterns of transfer in paired samples (**Figure 2**). A MASST search of these spectra against all publicly-available data in GNPS showed several matches to bacterial culture samples and gut/bile extracts, suggesting that these compounds may be of bacterial origin (MASST links in **Supplementary Table 2**).^17^ While the size of this dataset prevents a more comprehensive survey of which exogenous molecules can be directly transferred from the lactating parent to the infant via human milk, it does provide evidence that certain compounds can be taken up and metabolized by the infant.

**Figure 2:**
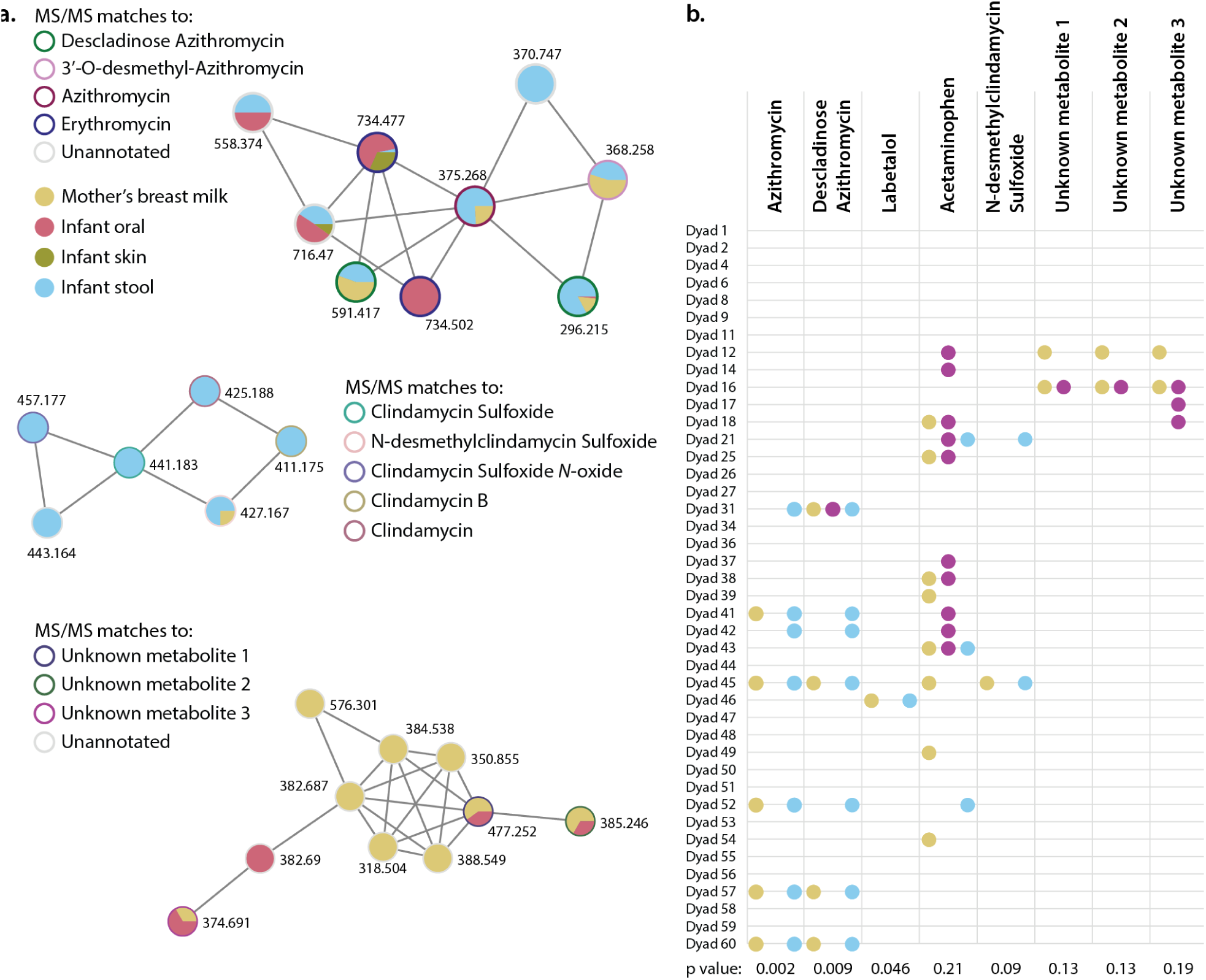
Drugs and their metabolites are transferred from the lactating parent’s milk to the infant. ***a***. Molecular networks of various drugs and other metabolites found in paired mother/infant dyads. Pie charts represent qualitative levels of antibiotics in each sampled area. ***b***. Appearance of paired compounds in each of the 40 mother/infant dyads tested. Colors correspond to type of sample. p values calculated by two-tailed Fisher’s Exact test.

Finally, we measured other exogenous metabolites in human milk, such as synthetic and food-derived compounds. Caffeine and caffeine metabolites dominated the list of food-derived compounds (**Supplementary Figure 2**), although compounds derived from black pepper, ginger, lentils, and cruciferous vegetables were also identified. A variety of plasticizers were also found in human milk, many of them common ingredients in personal care products. In fact, over 60% of the synthetic compounds identified by untargeted metabolomics are known cosmetic ingredients. This suggests that a metabolite does not have to be consumed orally by the lactating parent to affect the composition of human milk. Several of these exogenous metabolites were also identified in the paired samples. Food-derived compounds lenticin, theobromine, and 1,7-dimethyluric acid were identified in at least one dyad, along with the polyurethane precursor 1,5-naphthalenediamine.

## Discussion

While human milk is vital to infant health, we do not fully understand the scope of exogenous compounds (such as drugs or food components) that can be passed from the lactating parent to the infant via human milk. This is of critical importance since infant exposure to drugs such as antibiotics can have long-term negative effects on child health.^18–20^ Our analysis identified nine antibiotics along with an array of antibiotic metabolites in human milk, making it the largest and most metabolite-rich class of drugs identified. Several of the antibiotic metabolites did not match any known compounds in public databases, and even those compounds with annotations lacked critical information on biological activity. The variety of these molecules and the frequency with which they occur in human milk samples adds an additional layer of complexity to the transfer of antibiotics to the infant.

The data presented above suggest that exposure does not simply occur through ingestion by the infant, or even ingestion by the lactating parent, but may also be transferred by various routes into human milk. Our data suggest that a wide range of cosmetic compounds exist in human milk, as well as other topical synthetic compounds such as the insect repellent DEET (**Supplementary Table 1**). While it is possible that these topical compounds could have been ingested as contaminants from food products or entered milk via the nipple, the transfer of metabolites from the lactating parent’s skin to the infant via human milk is a novel mechanism that warrants further exploration. In addition, while untargeted metabolomics provides information on which compounds exist in human milk, it does not measure the exact concentrations of those components. Further pharmacokinetic studies are necessary to determine the safety of drugs or synthetic components in human milk, including exposure limits, half lives, and metabolism by both the host and microbes.^21^ However, our data does suggest that simply measuring the compound of interest in a pharmacokinetic study may not suffice. Further research is necessary to better describe the activity of the drug metabolites identified in this study in order to truly determine a drug’s safety in the infant.

Finally, while this study surveys ∼1000 subjects, it is still possible that sampling bias could affect our results. Thus, we do not present these data as a definitive list of exogenous compounds in human milk. Despite the gains in annotation achieved in this study, over 15,000 spectra remain unidentified (∼77% of those observed). This suggests that a rich diversity of metabolites in human milk remains unexplored.

---

**Supplementary Figure 1:**
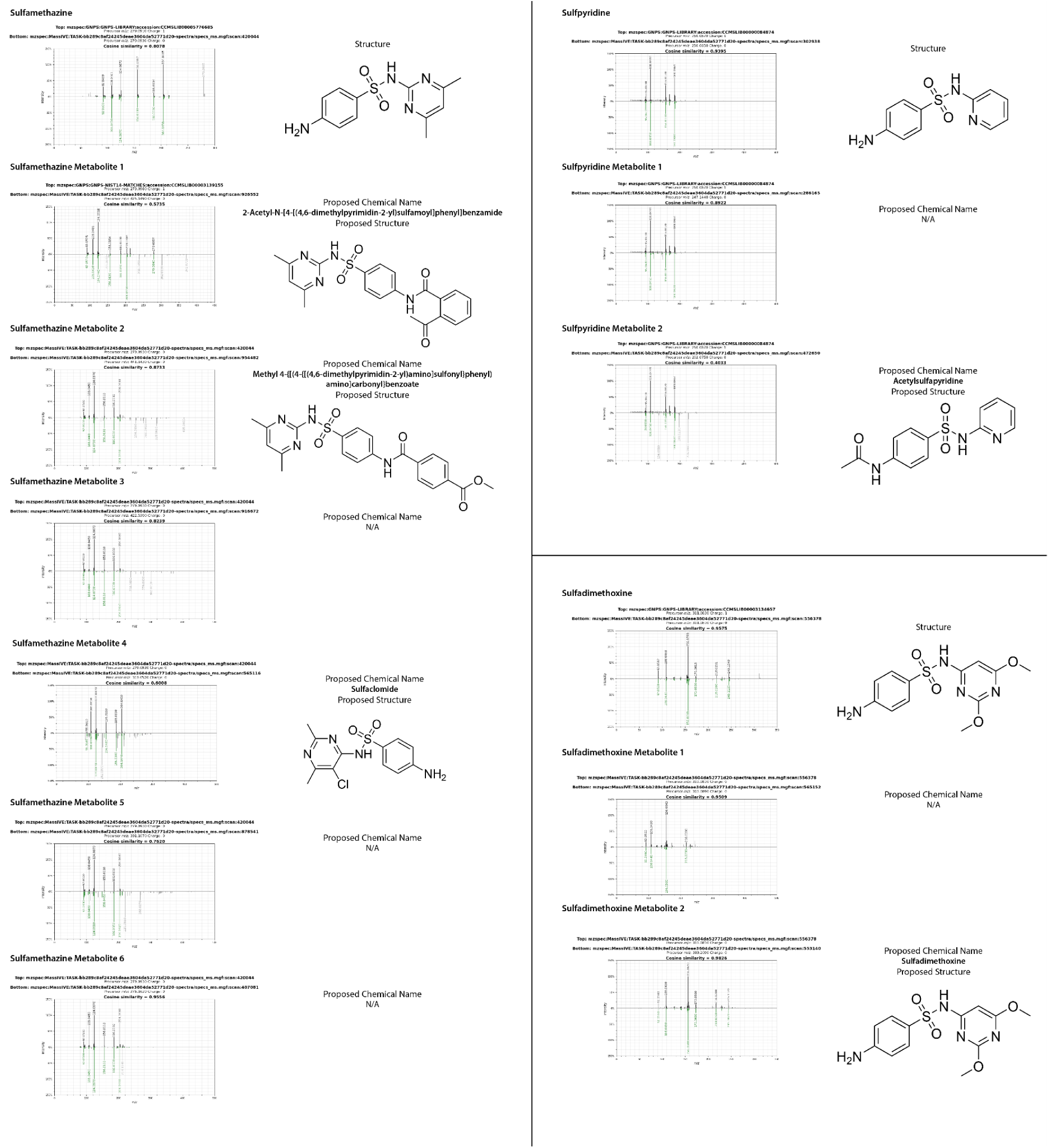
Sulfonamide antibiotic metabolites. Each metabolite has a mirror plot comparing a library/parent compound (top) and the antibiotic metabolite (bottom). Proposed chemical names and structures are listed when available.

## Code and data availability

Molecular networking jobs for each dataset (listed by accession numbers):

MSV000081571:

https://gnps.ucsd.edu/ProteoSAFe/status.jsp?task=7436a51b3749416699b7897c3194e9b6

MSV000081468:

https://gnps.ucsd.edu/ProteoSAFe/status.jsp?task=162f8a816c4e49489e58dcfa77a70e24

MSV000081432:

https://gnps.ucsd.edu/ProteoSAFe/status.jsp?task=e91e2e44e3234f08bb3d7f3f16d5f782

MSV000083463:

https://gnps.ucsd.edu/ProteoSAFe/status.jsp?task=071d91a251f7462a9a3d848c17980729

MSV000080074:

https://gnps.ucsd.edu/ProteoSAFe/status.jsp?task=6bcbb395727a48eeba599140697d4a2f

MSV000080117:

https://gnps.ucsd.edu/ProteoSAFe/status.jsp?task=0c31742f3fbb4c19b1c6821ec293ec0b

Breastmilk combined link:

https://gnps.ucsd.edu/ProteoSAFe/status.jsp?task=bb289c8af24245deae3604da52771d20

Breastmilk/stool/oral/skin combined link:

https://gnps.ucsd.edu/ProteoSAFe/status.jsp?task=680e3613b6b34092ad2f54a9ce552e5f

Passatutto FDR calculation:

https://gnps.ucsd.edu/ProteoSAFe/status.jsp?task=c2c8cc46b92d4725b8b0290a21448056

## Acknowledgements

We acknowledge funding from NIH grants T32HD123456 and P50HD106463.

## Notes

**Funding information:** T32HD123456, P50HD106463

https://gnps.ucsd.edu/ProteoSAFe/status.jsp?task=7436a51b3749416699b7897c3194e9b6

https://gnps.ucsd.edu/ProteoSAFe/status.jsp?task=162f8a816c4e49489e58dcfa77a70e24

https://gnps.ucsd.edu/ProteoSAFe/status.jsp?task=e91e2e44e3234f08bb3d7f3f16d5f782

https://gnps.ucsd.edu/ProteoSAFe/status.jsp?task=071d91a251f7462a9a3d848c17980729

https://gnps.ucsd.edu/ProteoSAFe/status.jsp?task=6bcbb395727a48eeba599140697d4a2f

https://gnps.ucsd.edu/ProteoSAFe/status.jsp?task=0c31742f3fbb4c19b1c6821ec293ec0b

https://gnps.ucsd.edu/ProteoSAFe/status.jsp?task=bb289c8af24245deae3604da52771d20

https://gnps.ucsd.edu/ProteoSAFe/status.jsp?task=680e3613b6b34092ad2f54a9ce552e5f

https://gnps.ucsd.edu/ProteoSAFe/status.jsp?task=c2c8cc46b92d4725b8b0290a21448056

### Competing Interest Statement

PCD is an advisor to Cybele and co-founder and scientific advisor of Ometa and Enveda with prior approval by UC-San Diego.

